# Proteolytic Activity and Substrate Specificity of Lake Geneva

**DOI:** 10.1101/2025.05.13.653673

**Authors:** Josephine Meibom, Natalie Wichmann, Michael Zumstein, Tamar Kohn

## Abstract

Heterotrophic microorganisms in lakewater secrete proteases which contribute to the turnover of dissolved organic matter and the degradation of peptidic contaminants. However, little is known about the identities and substrate specificities of these proteases. Herein, we sought to characterize the global proteolytic fingerprint of the extracellular proteases present in Lake Geneva, the largest freshwater body in Central Europe. Using Multiplex Substrate Profiling by Mass Spectrometry (MSP-MS), we identified preferred enzymatic cleavage next to positively charged and certain non-polar amino acids, while cleavage next to negatively charged residues was disfavored. Specifically, many of the detected cleavage sites were predominantly surrounded by arginine and lysine, consistent with a trypsin-like substrate specificity. This pattern was conserved across seasons and water depths and were shared with two other Swiss lakes. In contrast, we observed variability in the number and types of cleavage sites across samples, suggesting spatial and temporal differences in overall protease diversity. Using class-specific inhibitors, we found that serine and metalloproteases contribute to both exo- and endo-proteolytic activity in lakewater. Our findings expand our understanding of protein stability in lake ecosystems and may be used to predict the fate of peptidic contaminants in the environment.

## 1. Introduction

Dissolved organic matter (DOM) in lake ecosystems is a complex matrix comprised of organic molecules such as proteins, polysaccharides, and lipids^1,2^. Heterotrophic microorganisms can degrade DOM, thereby modulating its composition and turnover^1,3–5^. During this process, extracellular proteases are secreted to digest dissolved proteins into low molecular weight peptides and free amino acids which can be utilized by the microbial cells^4,6^. Besides DOM, these extracellular proteases may degrade peptidic contaminants such as antimicrobial peptides^7^ or viral pathogens^8^. The hydrolysis rates of antimicrobial peptides were found to be highly dependent on their amino acid sequences, indicating that the substrate specificities of the proteases present in environmental systems dictate the lifetimes of contaminants^7^. Similarly, bacterial proteases were found to cause the decay of human enteroviruses in lakewater^8^, though the antiviral effect was substrate specific^8,9^.

Protein degradation has been proposed to occur in two stages, where endoproteases first hydrolyze large proteins into smaller oligopeptides from which exoproteases subsequently release single amino acids^6^. In such a complex and dynamic system, individually identifying and characterizing all microbial extracellular proteases poses a major challenge. To date, the characterization of extracellular proteases in aquatic environments has focused on measurements of enzymatic activity levels^2,10–13^. These have often been performed using small fluorogenic substrates targeting leucine aminopeptidase (an exoprotease)^6^, resulting in a bias toward exo-rather than endo-proteolytic activity. Furthermore, the substrate specificities of the proteases present remain unexplored.

A suitable method to assess the substrate specificity of both exo- and endoproteases is Multiplex Substrate Profiling by Mass Spectrometry (MSP-MS)^14^. Employing a synthetic peptide library containing all possible peptide bonds, cleavage events are detected and used to generate a proteolytic fingerprint, representing the substrate specificity profile of the sample. The latter is visualized in the form of an iceLogo plot in which the amino acid specificity on either side of the cleavage site is displayed (**Figure 1**). Thus, the residues that are preferred or disfavored for cleavage by the proteases are identified. Although this method has been used to characterize the substrate specificities of microbial proteases produced by *Vibrio cholerae*^15^ and present in wastewater^16^, it has never before been applied to freshwater systems.

**Figure 1.**
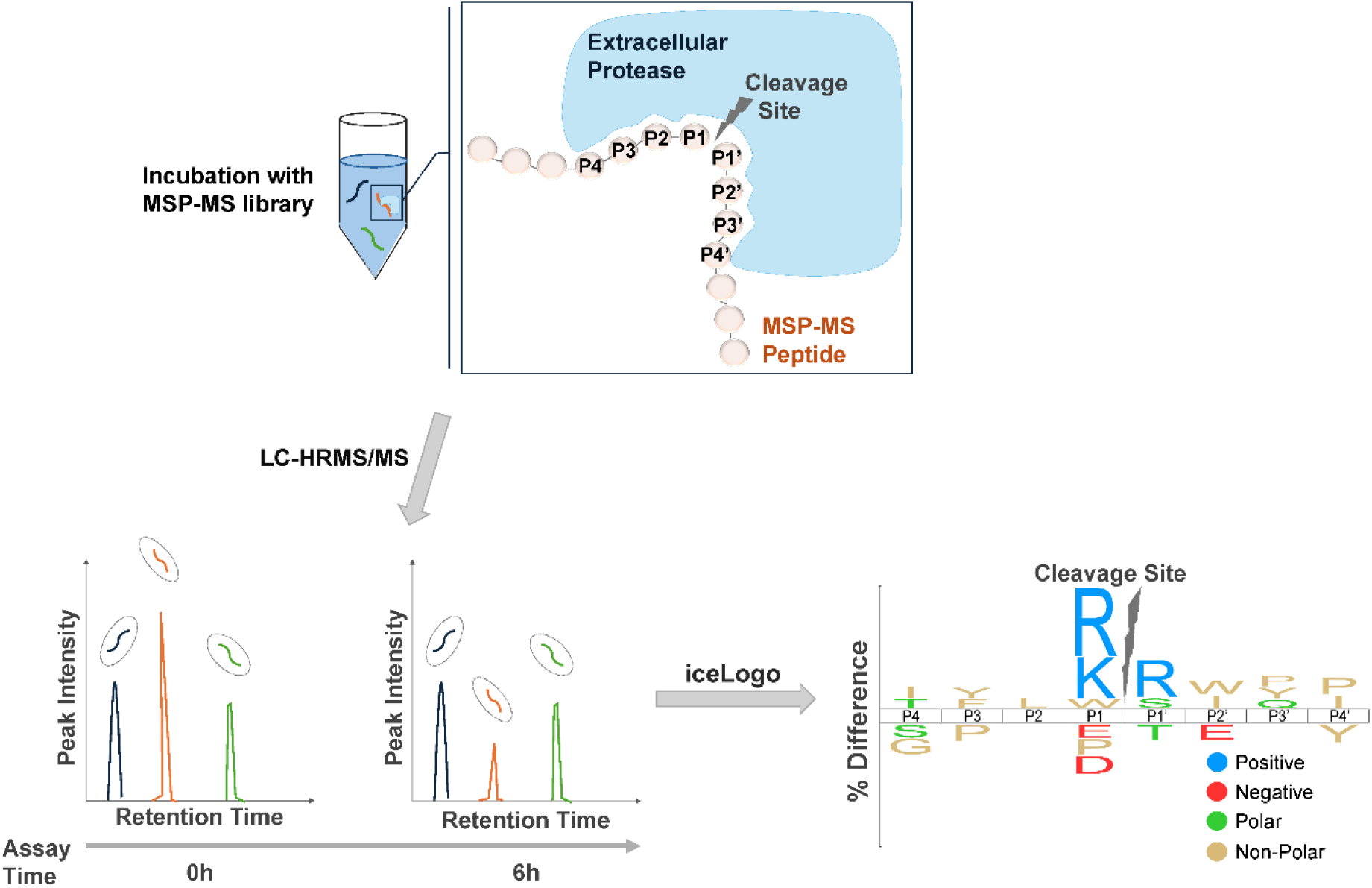
Workflow of the MSP-MS assay. Incubation of the MSP-MS library peptides in microbially active lakewater yields cleavage products which are detected by UHPLC-HRMS/MS. The eight residues (P4-P4’) surrounding the cleavage site are used to generate the proteolytic fingerprint of the sample in the form of an iceLogo plot.

Herein, we sought to describe the global substrate specificity profile of the microbial extracellular proteases present in Lake Geneva, the largest freshwater body in Central Europe. Using the MSP-MS assay, we characterized the proteolytic fingerprint of Lake Geneva as a function of seasonality and water depth and compared it to two other Swiss lakes of different trophic state. Together, these results provide insight into protein degradation in lake ecosystems and may further aid in understanding the fate of peptidic contaminants in this environment.

## 2. Materials and methods

### 2.1. Chemicals and Materials

Filter membranes with pore sizes of 0.8 μm (AAWP04700) and 0.22 μm MCE (GSWP04700) were purchased from MF-Millipore. MAXYMum Recovery microtubes of 1.5 mL (AXYMCT50LC) and 0.6 mL (AXYMCT060LC) volume were purchased from Sigma Aldrich. The DNeasy PowerWater Kit (14900-100-NF) was purchased from Qiagen and the Femto Bacterial DNA Quantification Kit (E2006) was purchased from Zymo Research.

Peptides of the MSP-MS peptide library^14^ were custom-synthesized by the University of Lausanne or by GenScript Biotech. 20 mM peptide stock solutions in dry DMSO (276855, Sigma Aldrich) were prepared in 0.6 mL MAXYMum Recovery microtubes. Each peptide was further diluted in 0.6 mL MAXYMum Recovery microtubes to yield 5 mM stock solutions in DMSO. 5 μL of each 5 mM stock solution were combined in a 1.5 mL MAXYMum Recovery microtube to yield the pooled MSP-MS peptide library containing all 124 peptides at a final concentration of 40.32 μM per peptide. Aliquots of the pooled peptide library were stored at -20 °C until use.

LCMS-grade acetonitrile (012041) was purchased from Biosolve. LCMS-grade formic acid (≥ 99%, 1EHK) was purchased from Carl Roth. GM6001 (CC1010), phenylmethanesulfonyl fluoride (PMSF, P7626), iodoacetamide (IAA, I6124), L-leucine-ß-naphthylamide hydrochloride (LLßN, L0376), ß-naphthylamine (ßN, 31618), and formaldehyde (≥ 36% in H2O, 47608) were purchased from Sigma Aldrich. SYTO 13 green fluorescent nucleic acid stain (5 mM solution in DMSO, S7575) was purchased from ThermoFisher Scientific. Working solutions of PMSF and ßN were prepared in analytical grade ethanol (≥ 99.8%, 10342652, Fisher Scientific). Working solutions of IAA and LLßN were prepared in deionized water.

### 2.2 Lakewater sampling

Lakewater samples were collected from Lake Geneva across seasons and depths, as well as from Lakes Neuchâtel and Bret. A map locating the three lakes and their sampling sites is presented in **Supplementary Figure 1** and the characteristics of each lake are listed in **Supplementary Table 1**. Surface lakewater was collected from Lake Geneva on April 9^th^ 2024, July 23^rd^ 2024, September 10^th^ 2024, and February 3^rd^ 2025 from the shore in Saint-Sulpice, Switzerland. On October 1^st^ 2024, lakewater was collected at depths of 1 m (surface), 34 m, and 65 m from the LéXPLORE platform (https://lexplore.info/) located 570 m from the shore of Pully, Switzerland. Surface lakewater was sampled from the shores of Lakes Neuchâtel (Yverdon-les-Bains, Switzlerand) and Bret on October 8^th^ 2024. Immediately after collection, the lakewater samples were brought to the laboratory and vacuum-filtered through a 0.8 μm MCE membrane filter to obtain microbially active lakewater, removing eukaryotes while retaining bacteria in the lakewater^17^. Half of the filtered lakewater was autoclaved at 121°C for 15 minutes to obtain sterile lakewater. All samples were stored at 4°C overnight prior to analysis of the proteolytic fingerprint (see section 2.4) and measurements of exo-proteolytic activity (see section 2.5).

### 2.3. Protease class inhibitor experiments

Lake Geneva water collected on September 10^th^ 2024 was used for the protease class inhibitor experiments. GM6001 was used as a metalloprotease inhibitor, phenylmethanesulfonyl fluoride (PMSF) was used as a serine protease inhibitor, and iodoacetamide (IAA) was used as a cysteine protease inhibitor. Each inhibitor was spiked individually into microbially active and sterile lakewater at a final concentration of 5 μM and let sit at room temperature for 30 min prior to analysis of the proteolytic fingerprint.

### 2.4. Determining the proteolytic fingerprint of lakewater by MSP-MS

The MSP-MS peptide library, designed by O’Donoghue *et al*^14^, is composed of 124 peptides of 14 residues each, giving rise to a total of 1612 possible cleavage sites. The central decapeptide regions host two copies of each amino acid pair (XY) and one copy of each X*Y and X**Y pair, with X and Y denoting specific amino acids and * representing randomly assigned amino acids. The central decapeptide acts as the substrate for endoproteases while distinct dipeptides are present at the termini to allow the identification of exo-proteolytic specificities. Thus, both exo- and endo-proteolytic cleavage events are accounted for and used to generate the proteolytic fingerprint. Cysteine was not included in the MSP-MS library to avoid disulfide bond formation, and oxidation-prone methionine was replaced by its isosteric analog norleucine (indicated as M). The prevalence of each amino within the MSP-MS library spans 4.2% to 6.8%.

**Figure 1** outlines the workflow of the MSP-MS assay. Microbially active and sterile lakewater samples were brought up to room temperature prior to incubation with the MSP-MS peptide library. The MSP-MS protocol devised by O’Donoghue *et al*.^14,18^ was followed with an adapted enzyme quenching process^16^. Specifically, the peptide library was spiked into 500 μL microbially active or sterile lakewater (in triplicate) in 0.6 mL or 1.5 mL MAXYMum Recovers microtubes to a final peptide concentration of 0.5 μM per peptide. At different timepoints, 55 μL of solution were transferred into a new tube and heat treated at 80°C (water bath) for 10 minutes to inactivate the proteases. The inactivated samples were centrifuged for 1 minute at maximum speed (25’000 *g*) and the supernatant (50 μL) was transferred into a new tube. The samples were stored at -20°C until further analysis by liquid chromatography coupled to high-resolution mass spectrometry.

The MSP-MS library peptides and their cleavage products were analyzed by Ultra-High-Pressure Liquid Chromatography coupled to High-Resolution Tandem Mass Spectrometry (UHPLC-HRMS/MS) using a Vanquish Flex UHPLC (ThermoFisher Scientific) equipped with an ACQUITY UPLC HSS T3 Column (REF#: 186003539) and coupled to an Orbitrap Exploris 120 (ThermoFisher Scientific). A sample volume of 10 μL was injected (1 μg total peptide) at a flow rate of 300 μL/min and using the following elution gradient (A: 0.1% LCMS-grade formic acid in ultrapure water, B: 0.1% LCMS-grade formic acid in LCMS-grade acetonitrile): 0-16 min: 5% B – 30% B, 16-18 min: 30% B – 90% B, 18-19 min: 90% B, 19-19.5 min: 90% B – 5% B, 19.5-22 min: 5% B. Detection parameters were set to: MS full-scan: range: 150 – 2000 m/z, resolution: 120’000, AGC target: Standard, Maximum IT: Auto, positive electrospray ionization (tune data: ion transfer tube temperature: 360 °C, sheath gas: 40, aux gas: 15, sweep gas: 1, RF Lens: 70.0), MS/MS acquisitions: resolution: 15’000, AGC target: Standard, Maximum IT: Auto, dynamic exclusion time: 60 ms. Skyline (version 24.1.0.199) was used for parent peptide degradation analysis. The following criteria for parent peptides and hydrolysis product identification were used: m/z deviation < 2 ppm, MS2 fragments with m/z deviation < 5 ppm.

Data analysis was performed in R (version 4.4.3) according to the procedure described by Rohweder *et. al*^18^. Peaks Studio 11 (Bioinformatics Solutions Inc.) was used to perform label-free quantification (LFQ) and database search against the MSP-MS library peptide sequences. Precursor tolerance was set to 20 ppm and 0.01 Da for MS2 fragments. No protease digestion was specified and a peptide length of 3-14 amino acids was defined. No normalization was used for LFQ. A false discovery rate (FDR) threshold was set to < 1 % for all peptides. R scripts kindly provided by the O’Donoghue laboratory (University of California San Diego, link: https://github.com/briannahurysz/MSP-MS) were used to filter out peptides with a quality score ≤ 0.3. NormalyzerDE^19^ was utilized for data evaluation and normalization. The R scripts were used to identify the eight residues (P4-P4’) surrounding the cleavage site. Only cleavage products with intensity scores eightfold or higher than the intensity scores in the t = 0h sample were considered significant. Moreover, Dixon’s Q test was implemented to eliminate outliers from replicate samples. Timepoints (6h or 24h) were selected for sample comparison to ensure that all samples provided enough cleavage events to generate representative iceLogo plots, P1↓P1’ cleavage Venn diagrams, and P1↓P1’ cleavage frequency heatmaps.

Using the identified P4-P4’ sequences (positive set), the iceLogo software^20^ was used to generate the proteolytic fingerprint of the sample. The percent difference between the positive and reference (MSP-MS peptide library) sets of the frequency of an amino acid at each P4-P4’ position was calculated. Only significantly (t-test, P ≤ 0.05) under- or over-represented residues at each position are included in the iceLogo plots. The identified P4-P4’ sequences were additionally used to generate P1↓P1’ cleavage Venn diagrams and frequency heatmaps (↓ indicates the cleavage site). The Venn diagrams were made using the identified P1↓P1’ pairs in each sample and the VennDiagram R package. For the P1↓P1’ cleavage frequency heatmaps, the occurrences of each cleaved P1↓P1’ pair were divided by their respective occurrences in the MSP-MS peptide library, yielding the cleavage frequency (%) of that pair. The heatmaps were generated using the ggplot2 R package. Finally, pairwise comparisons of the amino acid frequencies at positions P4-P4’ in samples April 2024, July 2024, September 2024, October 2024, and February 2025 were made by calculating Pearson correlation coefficients. The Benjamini-Hochberg method was used to adjust the calculated p-values.

### 2.5. Exo-proteolytic activity measurement

Hydrolysis of the substrate L-leucine-ß-naphthylamide hydrochloride (LLßN) was used to measure the exo-proteolytic activity exerted by leucine aminopeptidase and similar aminopeptidases in microbially active lakewater, based on a method previously described^21^. In brief, 100 μL of LLßN solution (2 mM) was mixed with 100 μL sample in a well of a 96 well-plate to achieve an initial substrate concentration of 1 mM. Each sample was prepared in triplicate and was brought to room temperature prior to incubation with the LLßN substrate. The reactions were incubated for 4h in the dark at room temperature. Subsequently, fluorescence of ß-naphthylamine (ßN) was measured with 340 nm excitation and 410 nm emission using a Biotek Synergy MX microplate reader. The same procedure was performed for sterile lakewater. The average fluorescence intensity of the sterile lakewater was subtracted from that of the microbially active lakewater. The fluorescence intensity was converted into concentration of ßN using a calibration curve of ßN standards. Exo-proteolytic activity was expressed as the concentration of ßN [μM] produced during the 4h incubation period.

### 2.6. Quantification of bacteria in lakewater by flow cytometry

Bacteria in the microbially active lakewater samples of Lake Geneva (October 1^st^ 2024) and lakes Neuchâtel and Bret (October 8^th^ 2024) were quantified by flow cytometry. Each sample was counted in triplicate. In brief, bacterial cells in the samples were fixed by mixing 900 μL sample with 100 μL 36% formaldehyde. The fixed samples were stored at 4°C until analysis. Immediately prior to analysis, the bacterial cells were stained by adding 2 μL SYTO 13 nucleic acid stain to 198 μL fixed sample. The mixture was incubated at 37°C for 5 minutes, followed by 15 minutes in the dark at room temperature. 50 uL were injected onto an Attune CytPix volumetric flow cytometer (ThermoFisher Scientific). The bacterial cells were identified using green fluorescence and forward scatter properties. Unstained samples were used as controls.

## 3. Results

### 3.1. The proteolytic profile of Lake Geneva throughout a year

To characterize the proteolytic profile of Lake Geneva across different seasons, we first measured exo-proteolytic activity in lakewater collected between April 2024 and February 2025 using LLßN as a fluorogenic substrate. We observed highly variable levels of exo-proteolytic activity (**Figure 2A**) with April 2024 presenting 5-to 40-fold higher levels of activity compared to all other samples. Next, the MSP-MS method was employed to determine the proteolytic fingerprint of the same samples. We monitored the number of library peptide cleavage sites throughout the incubation period and observed a continuous increase in cleavage site numbers over 24h for all samples (**Figure 2B**). Although April 2024 presented the highest level of exo-proteolytic activity and the highest number of cleavage sites as detected by MSP-MS, the number of cleavage sites did not globally follow the same trend as for exo-proteolytic activity across all samples (**Supplementary Figure 2**).

**Figure 2.**
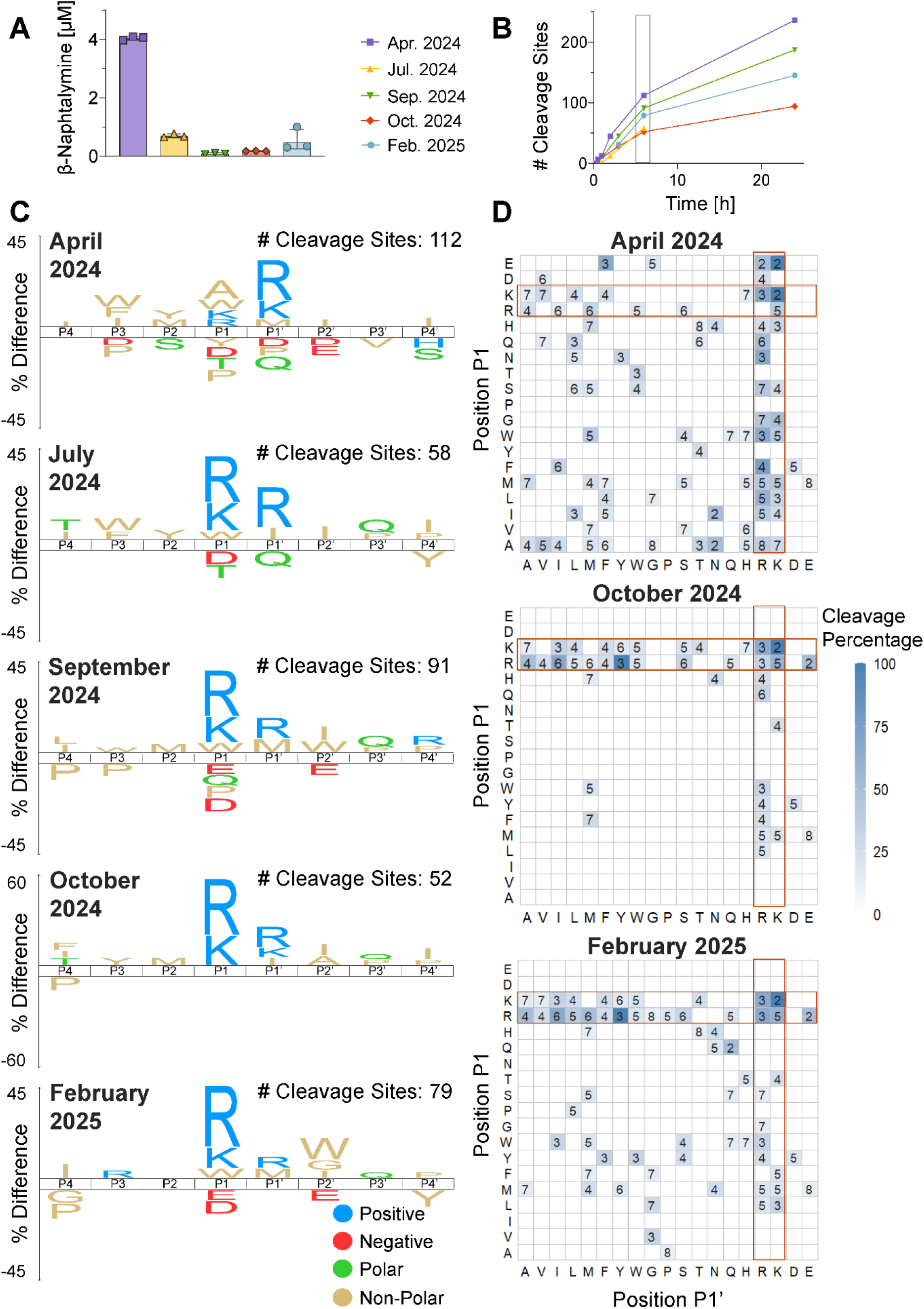
Characterization of the proteolytic profile of Lake Geneva throughout a year. **(A)** Exo-proteolytic activity at t = 6h measured by the LLßN assay. Individual data points of triplicate measurements are presented with the geometric mean and standard deviation. **(B)** The number of detected cleavage sites during incubation of the MSP-MS peptide library in Lake Geneva water. Note that no measurement was made at t = 24h for July 2024. The grey box indicates t = 6h, for which time iceLogo plots and P1↓P1' cleavage frequency heatmaps are presented in **(C)** and **(D)** respectively. **(C)** Proteolytic fingerprints (t = 6h) of Lake Geneva, presented as iceLogo plots. The number of detected cleavage sites at t = 6h is indicated for each sample. One-letter amino acid codes are used with M representing norleucine. Note the change in scale for October 2024. **(D)** P1↓P1' cleavage frequency heatmaps at t = 6h for April 2024, October 2024, and February 2025. The number in each cell represents the number of occurrences of the corresponding P1↓P1' pair in the MSP-MS library. The color scale indicates the percentage of the different P1↓P1' pairs that are cleaved during incubation in the sample. Horizontal and vertical orange boxes highlight the main amino acid specificities (K & R) at positions P1 and P1' respectively.

Using the detected cleavage sites for each sample, we generated the proteolytic fingerprints of Lake Geneva in the form of iceLogo plots. We observed a change in the proteolytic fingerprints throughout the incubation period (**Supplementary Figure 3**) in accordance with the increasing number of cleavage sites (**Figure 2B**). An incubation time of 6h was selected to compare samples, as sufficient cleavage events were observed in all samples at this timepoint. Across seasons, we observed a preference for positively charged (arginine (R) and lysine (K)) amino acids around the cleavage site (P1↓P1’) (**Figure 2C**), while some non-polar (tryptophan (W), norleucine (M), and isoleucine (I)) residues were also commonly observed. Certain aspects of these patterns were similar to the serine proteases trypsin and chymotrypsin, which favor cleavage of arginine and lysine and of aromatic residues at position P1 respectively^22^. Interestingly, the specificity at position P1 was dominant for all samples except for April 2024, for which the specificity at position P1’ was prevalent. Our results further revealed a distinct disfavoring of negatively charged (glutamic acid (E) and aspartic acid (D)) and polar (glutamine (Q), threonine (T), and sometimes serine (S)) residues, with the exception of glutamine which was occasionally preferred at position P3’.

To fully analyze the proteolytic fingerprint of each sample, we plotted the cleavage frequencies of all P1↓P1’ pairs as a heatmap (**Figure 2D, Supplementary Figure 4**). This confirmed the dominant preference for arginine and lysine at positions P1 and P1’ (**Figure 2D**, orange bands) observed previously in the iceLogo plots. Moreover, the distinct importance of position P1’ (orange vertical band) for substrate specificity in April 2024 was also emphasized. Calculation of pairwise Pearson correlation coefficients of the amino acid frequencies at positions P4-P4’ (**Supplementary Table 2**) further validated that April 2024 was the most unique of all samples analyzed.

Despite the broad similarity of the proteolytic fingerprints, the P1↓P1’ cleavage frequency heatmaps (**Figure 2D, Supplementary Figure 4**) revealed the presence of additional minor (i.e. those not belonging to the R/K bands) proteolytic substrate specificities. While some of these, such as H↓M, W↓M, W↓S, F↓M, and M↓E, were common to all or most samples, many were unique or only shared by a few samples. For example, April 2024 was the only sample to possess some specificity towards E and D as well as high specificity towards A at position P1. To further evaluate these differences, we determined how many cleaved P1↓P1’ pairs were shared among the samples (**Supplementary Figure 5**). While sixteen pairs were common to all samples, many were cleaved only in two or three. A large variability in the number of unique P1↓P1’ pairs per sample was observed, with April 2024 presenting thirty-four unique cleaved P1↓P1’ pairs and October 2024 possessing none. Overall, these results confirm that April 2024 possessed many minor substrate specificities and indicate that the diversity of cleavage sites varies across samples.

### 3.2. Proteolytic profiles of Lake Geneva at different depths

We next studied the effect of depth on the proteolytic profile of Lake Geneva. First, we measured the temperature profile throughout the water in October 2024. We observed thermal stratification (**Figure 3A**) with a maximal temperature of 15°C at the surface and a minimal temperature of 8°C at depths greater than 50 m. Based on these temperatures, we selected three depths for further analysis: 1 m (15°C, epilimnion), 34 m (12°C, thermocline), and 65 m (8°C, hypolimnion). Next, we measured exo-proteolytic activity and counted bacterial cell numbers at each depth (**Figure 3B**), revealing that both parameters decreased with increasing depth.

**Figure 3.**
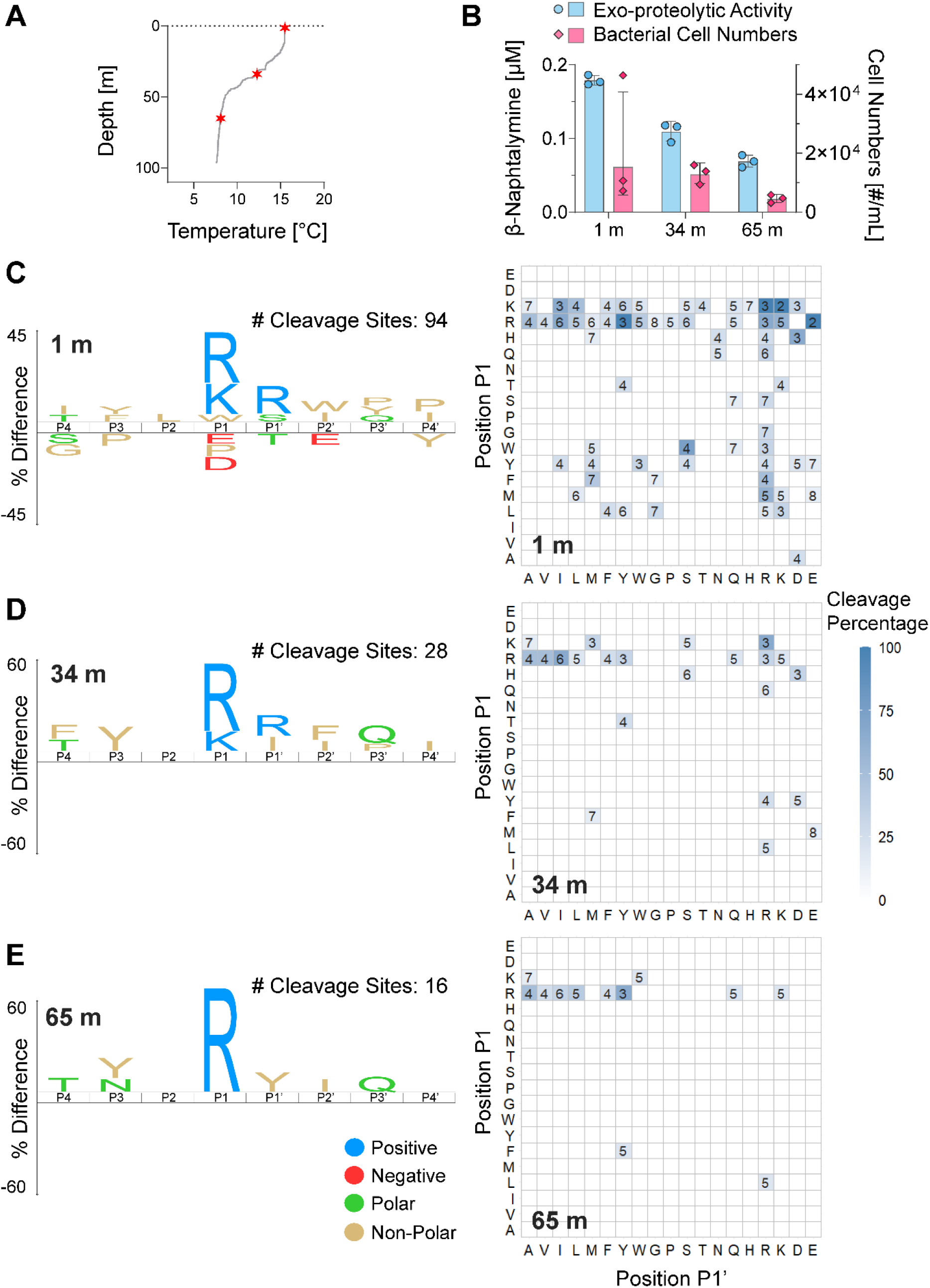
Characterization of the proteolytic profile of Lake Geneva at different depths. **(A)** Temperature profile of Lake Geneva (October 2024). From this, three depths (indicated by red stars) were selected for further analysis: 1 m, 34 m, and 65 m. **(B)** Exo-proteolytic activity as measured by the LLßN assay and bacterial cell numbers determined by flow cytometry at the three selected depths. Individual data points of triplicate measurements are presented with the geometric mean and standard deviation. (C-E) Proteolytic fingerprints presented as iceLogo plots (note the different scales), and P1↓P1' cleavage frequency heatmaps of Lake Geneva at the selected depths (t = 24h): 1 m (C), 34 m (D), and 65 m (E).

Incubation with the MSP-MS peptide library for 24h allowed us to generate the proteolytic fingerprint at each depth in the form of iceLogo plots and P1↓P1’ cleavage frequency heatmaps (**Figure 3C-E**). It should be noted that all assays were performed at room temperature and not at the original (lower) lakewater temperature. Performing the MSP-MS assay at the original lakewater temperature led to fewer detected cleavage sites over the 6h incubation period considered here (**Supplementary Figure 6**). The MSP-MS method thus represents the sample’s proteolytic potential and does not fully mimic the true environmental conditions at each depth. We observed decreasing numbers of cleavage sites and slower peptide cleavage kinetics with depth (**Supplementary Figure 7**), following the trends in exo-proteolytic activity and bacterial cell numbers. In accordance with the lower number of cleavage sites, the iceLogo plots and cleavage frequency heatmaps at 34 m and 65 m depth (**Figure 3D&E**) appeared less populated than those at 1 m. In deeper lakewater, the cleavage specificity was dominated by arginine and lysine mainly at position P1 while the specificity for these residues at position P1’ was either lost or did not manifest over the incubation period. Overall, there was a loss in diversity of the proteolytic fingerprint with water depth.

### 3.3. Identification of main extracellular protease classes present in Lake Geneva

To determine the catalytic classes of the extracellular proteases present in Lake Geneva, we employed class-specific protease inhibitors. We measured the exo-proteolytic activity of Lake Geneva in the presence of different inhibitors targeting cysteine, serine, metalloproteases (**Figure 4A**). Unexpectedly, the addition of the cysteine protease inhibitor led to an increase in exo-proteolytic activity, though the cause for this increase is unknown. In contrast, we observed a complete absence of exo-proteolytic activity in the presence of inhibitors targeting serine and metalloproteases. This finding may be partly rationalized by the effect of inhibitors on the activation processes required by some extracellular proteases upon secretion, sometimes performed by proteases of another class. For example, the metalloprotease staphylolysin produced by *Pseudomonas aeruginosa* is activated extracellularly upon cleavage of its propeptide by pseudolysin (a metalloprotease) or by lysine-specific endopeptidase (a serine protease), two proteases also secreted by this bacterium^23^. The presence of either metalloprotease or serine protease inhibitors would thus result in the suppression of staphylolsin activity.

**Figure 4.**
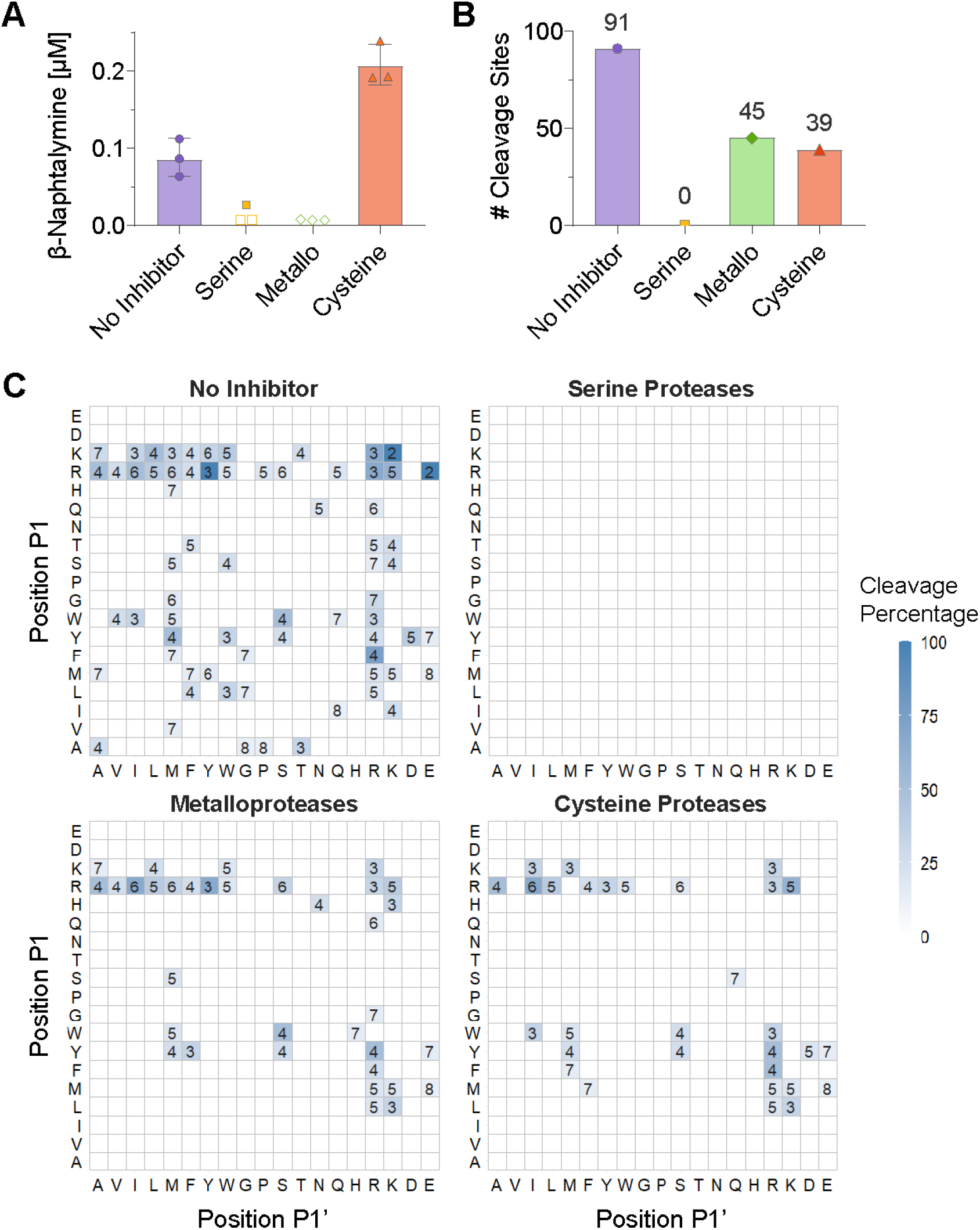
The effect of inhibitors on the proteolytic profile of Lake Geneva (September 2024). Inhibitors targeting one protease class were added separately to microbially active lakewater at a final concentration of 5 μM. Phenylmethanesulfonyl fluoride (PMSF) was used as the serine protease inhibitor, GM6001 was used as the metalloprotease inhibitor, and iodoacetamide (IAA) was used as the cysteine protease inhibitor. **(A)** Exo-proteolytic activity as measured by the LLßN in the absence or presence of class-specific protease inhibitors. Individual data points of triplicate measurements are presented with the geometric mean and standard deviation. Open symbols represent values below the detection limit. **(B)** The number of detected MSP-MS library peptide cleavage sites in the absence or presence of class-specific protease inhibitors (t = 6h). **(C)** P1↓P1’ cleavage frequency heatmaps (t = 6h).

Next, the proteolytic fingerprint of each sample was determined using the MSP-MS method. In control experiments with sterile lakewater, we confirmed that the inhibitors did not interfere with the stability of the library peptides as no significant loss of peptide occurred over 6h incubation time (**Supplementary Figure 8**). We observed zero cleaved P1↓P1’ pairs in the presence of the serine protease inhibitor and an approximately two-fold reduction in the number of detected cleavage sites in the presence of both the cysteine and metalloprotease inhibitors (**Figure 4B**). The addition of cysteine and metalloprotease inhibitors partially removed cleavage specificity associated with arginine and lysine at positions P1 and P1’ as well as some minor P1↓P1’ pairs (**Figure 4C**). The similarity of the P1↓P1’ cleavage frequency heatmaps in the presence of these inhibitors may be a result of the interdependence of proteases of different classes, as discussed above. However, these results also suggest that these two protease classes may be associated with similar proteolytic cleavage specificities in Lake Geneva.

### 3.4. Comparison of the proteolytic profile of three Swiss lakes

To understand whether the proteolytic profile of Lake Geneva was unique or shared among other freshwater sources in Switzerland, we compared mesotrophic Lake Geneva with two other Swiss lakes of different trophic state (Lakes Neuchâtel (oligotrophic) and Bret (eutrophic)) in October 2024. A map locating the three lakes and their sampling sites is presented in **Supplementary Figure 1** and the characteristics of each lake are listed in **Supplementary Table 1**. The three lakes presented different levels of exo-proteolytic activity and bacterial cell numbers (**Figure 5A**), with Lake Bret featuring the highest levels of both parameters and Lake Geneva the lowest. The MSP-MS method revealed differences in peptide cleavage kinetics (**Figure 5B**). We observed a steep initial increase in the number of cleavage sites in Lakes Geneva and Bret followed by a slower rise from 6h to 24h incubation time while a linear increase was observed in Lake Neuchâtel throughout the incubation period, suggesting differences in enzyme kinetics in each water source. At 6h incubation time, Lake Bret had a larger number of cleavage sites compared to Lakes Geneva and Neuchâtel, which had similar numbers. Despite these differences, the dominant features of the proteolytic fingerprint of Lakes Neuchâtel and Bret (i.e., a preference for positively charged amino acids and a disfavoring of negatively charged residues around the cleavage site, **Supplementary Figure 9**) were similar to that described in section 3.1. for Lake Geneva (**Figure 2C**).

**Figure 5.**
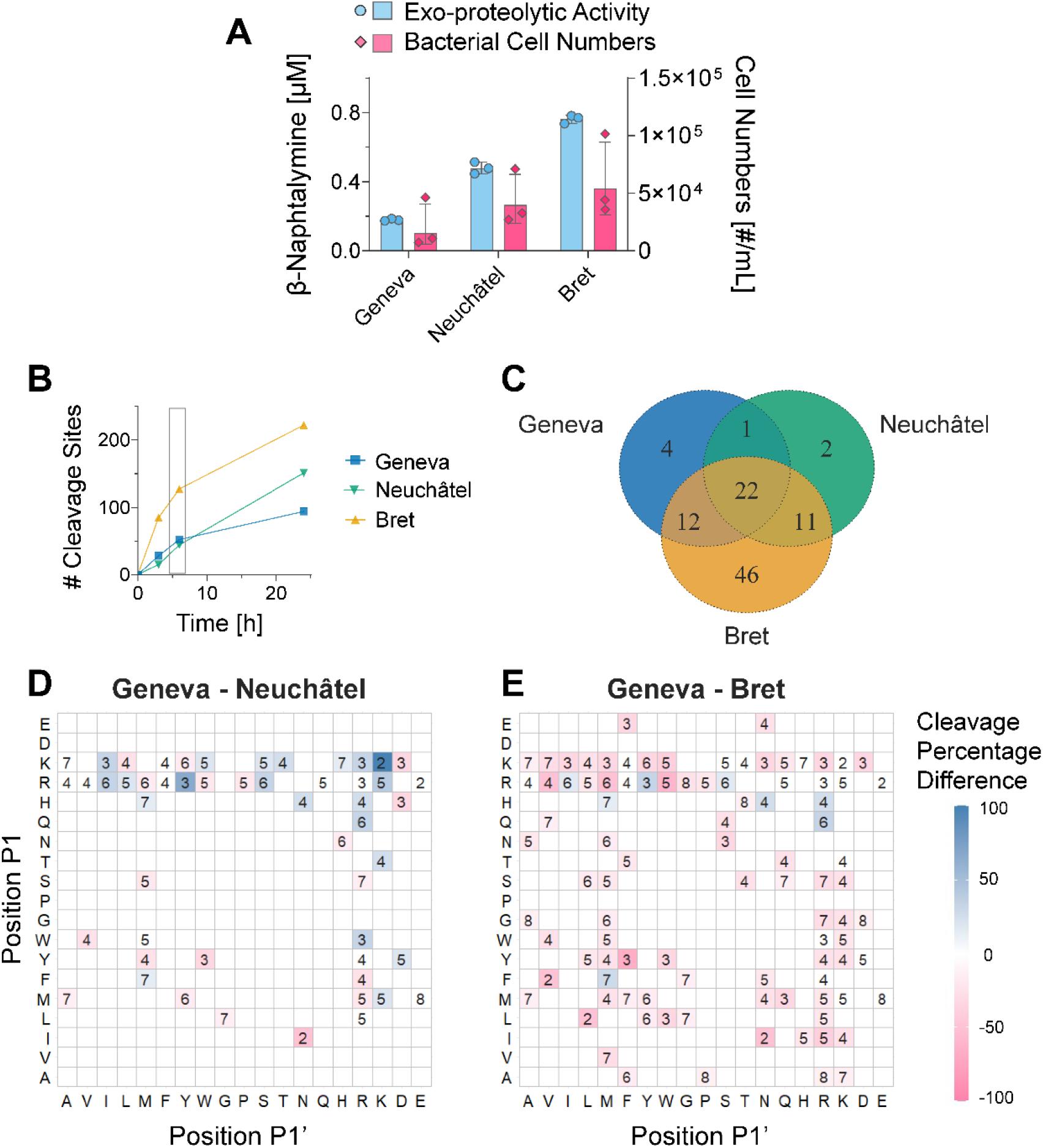
Characterization of the proteolytic profile of three different Swiss Lakes. **(A)** Exo-proteolytic activity as measured by the LLßN assay and bacterial cell numbers determined by flow cytometry in Lakes Geneva, Neuchâtel, and Bret measured in October 2024. Individual data points of triplicate measurements are presented with the geometric mean and standard deviation. **(B)** The number of detected cleavage sites during incubation of the MSP-MS peptide library in lakewater. The grey box indicates t = 6h, for which time P1↓P1’ cleavage frequency difference heatmaps are presented in (C) and (D). **(C)** Venn diagram representing the different P1↓P1’ pairs cleaved in each lake (t = 6h). Each cleaved P1↓P1’ pair is counted once, irrespective of its cleavage frequency in the sample. **(D-E)** P1↓P1’ cleavage frequency difference heatmaps (t = 6h). The cleavage frequencies of one lake are subtracted from another to yield the difference heatmap: **(D)** Lake Geneva minus Lake Neuchâtel, **(E)** Lake Geneva minus Lake Bret. Cells colored in blue correspond to P1↓P1’ pairs that are dominantly cleaved in Lake Geneva, while cells colored in pink correspond to pairs that are dominantly cleaved in the second lake. P1↓P1’ pairs cleaved equally in both lakes are indicated by white cells containing the number of occurrences of the pair in the MSP-MS library.

To evaluate the differences in their proteolytic fingerprints, we counted how many cleaved P1↓P1’ pairs were shared among the three lakes (**Figure 5C**). While most of the P1↓P1’ pairs cleaved in Lakes Geneva and Neuchâtel were also cleaved in one or both of the other lakes, almost half of the cleaved pairs present in Lake Bret were unique to this water source. Next, we plotted P1↓P1’ cleavage frequency difference heatmaps (**Figure 5D-E**) in which the cleavage frequency of each P1↓P1’ pair in Lake Neuchâtel (**Figure 5D**) or Lake Bret (**Figure 5E**) was subtracted from its frequency measured in Lake Geneva. We observed equal cleavage frequencies for twelve P1↓P1’ pairs in Lakes Geneva and Neuchâtel (**Figure 5D**), while Lake Geneva presented higher cleavage frequencies compared to Lake Neuchâtel for pairs involving arginine and lysine at positions P1 and P1’. Minor P1↓P1’ pairs (not belonging to the R/K bands) were predominantly cleaved in Lake Neuchâtel. We observed equal cleavage frequencies for eighteen P1↓P1’ pairs in Lakes Geneva and Bret and higher cleavage frequencies in Lake Bret for almost all other P1↓P1’ pairs (**Figure 5E**). This reflected the much higher number of cleavage sites observed in Lake Bret at 6h compared to Lake Geneva (**Figure 5B**) as well as the many unique P1↓P1’ pairs present in Lake Bret (**Figure 5C**), indicating a high diversity of extracellular proteolytic substrate specificity in this lake.

## 4. Discussion

In this study, we determined the proteolytic fingerprint of Lake Geneva thereby revealing the overall substrate specificity of the extracellular proteases present in the largest lake in Central Europe. We found that positively charged amino acids, as well as certain non-polar residues, were preferred on either side of the cleavage site, while negatively charged and polar amino acids were not. These patterns were conserved throughout the year and at different depths and were also present in two other Swiss lakes.

The high conservation of the prominent features of the proteolytic fingerprint of Lake Geneva throughout the year (**Figure 2C&D**) suggests either that certain microbial extracellular proteases dominate throughout time or that many different proteases possess similar substrate specificities. The dominant preference for arginine and lysine residues at positions P1 and P1’ revealed “trypsin-like” activity in lakewater while the presence of tryptophan pointed toward “chymotrypsin-like” specificity^22^. Interestingly, similar proteolytic fingerprints have been reported in municipal wastewater^16^ suggesting conserved global substrate specificities for microbial extracellular proteases across different freshwater environments. Importantly, highly dissimilar profiles were reported for two proteases involved in colony formation by *Vibrio cholerae*^15^, confirming that the similarities noted above are not attributable to any bias in the MSP-MS assay.

While the main patterns observed in the proteolytic fingerprints were conserved, exo-proteolytic activity levels and the diversity of cleaved P1↓P1’ pairs were highly variable (**Figure 2A, Supplementary Figure 5**). In agreement with our results, daily and seasonal variations in exo-proteolytic activity have been reported in marine^10,24^ and lake^11,25^ environments. Such differences may be due to seasonal variations in the relative abundances of the extracellular proteases or by the periodic production of rare proteases. Factors that influence the secretion of extracellular proteases include the concentration^6,2,25–29^ and identity^30,31^ of substrates present in the environment. In general, protease production has been found to be repressed by readily-assimilable nutrients and to be stimulated in the presence of larger oligopeptides found in DOM^6,26,28^. Thus, variations in DOM composition across seasons^1,3^ and with rainfall^5^ may account for the differences observed here. Additionally, specific substrates can directly promote protease secretion. For example, certain tripeptides were found to induce the expression of the extracellular protease MCP-01 in *Pesudoalteromonas* sp. SM9913, a bacterium isolated from deep-sea sediments^31^. Based on these findings, peptides present in the MSP-MS library could potentially induce the secretion of proteases during the incubation period and thereby influence the proteolytic fingerprints. The short incubation times used here (6h or 24h) should however minimize such an effect^32^. The diversity of cleaved P1↓P1’ pairs and the variability in the number of unique pairs across samples (**Supplementary Figures 4&5**) suggest that minor proteolytic substrate specificities (i.e., those not belonging to the dominant R/K bands) may be associated with extracellular proteases produced by periodically present (rare) bacteria. In agreement with this suggestion, fluctuations of the relative abundances of bacterial phyla have been observed in Lake Geneva across seasons, with significant variations of minor bacterial species^33^. Thus, seasonal factors such as variations in DOM and bacterial community composition may significantly affect proteolytic substrate specificities associated with rare extracellular proteases, while the global extracellular proteolytic fingerprint is conserved.

These considerations can also be applied to our study of different depths in Lake Geneva, in which we observed a decrease in bacterial cell numbers, exo-proteolytic activity, and MSP-MS library peptide cleavage sites with increasing depth (**Figure 3**). Changes in bacterial populations with water depth have been linked to variations in exo-proteolytic activity in sea^29,34^ and lakewater^11^. In these studies, correlations have been found between exo-proteolytic activity and bacterial numbers^29,34^, in agreement with our results (**Supplementary Figure 2C**). The lower numbers of MSP-MS cleavage sites detected in deeper waters were consistent with the lower exo-proteolytic activities and may be related to differences in bacterial numbers and community composition at each depth. Indeed, substantial differences in the bacterial communities in the top (0-30 m depth) and bottom (30-100 m depth) layers of Lake Geneva have been reported^33^. Although the dominant preference for arginine and lysine at positions P1 and P1’ was retained, a clear loss in proteolytic substrate diversity was observed (**Figure 3C-E**) at lower depths.

The three lakes studied here presented different levels of exo-proteolytic activity, bacterial cell numbers, and MSP-MS library peptide cleavage sites (**Figure 5A&B**). As reported in other studies, these results reflect the different trophic states of the lakes^1–3,35^, classified as mesotrophic^36^, oligotrophic^37^, and eutrophic^38^ for Lakes Geneva, Neuchâtel, and Bret respectively. The many unique P1↓P1’ cleavage pairs detected in Lake Bret reflect its high biological activity and are likely indicative of a wider variety of proteinaceous substrates present in this water source. However, the main features of the proteolytic fingerprint of Lake Geneva were also present in Lakes Neuchâtel and Bret (**Supplementary Figure 9**), suggesting a common set of microbial extracellular proteases or proteases with similar substrate specificities in all three lakes.

Consistent with other studies^8,11,13,27^, we found that the exo-proteolytic activity of Lake Geneva was fully suppressed in the presence of serine and metalloprotease inhibitors (**Figure 4A**), suggesting that these two protease classes are predominant in lakewater. Interestingly, we observed full suppression of MSP-MS library peptide cleavage events in the presence of the serine protease inhibitor but only a two-fold reduction in the presence of the metalloprotease inhibitor (**Figure 4B**). Furthermore, the low number of MSP-MS cleavage sites did not agree with the high exo-proteolytic activity measured in the presence of the cysteine protease inhibitor. The discrepant effects of inhibitors on exo-proteolytic activity and proteolytic fingerprints may be partially explained by the differences in what the two assays measure. While the proteolytic fingerprint represents the activity of both exo- and endo-proteases, the exo-proteolytic assay only captures a small subset of proteases. Additionally, the inhibition of one class may affect the activity of proteases of another, as some proteases require activation upon secretion.

Surprisingly, the addition of class-specific protease inhibitors to lakewater did not reveal large variations in proteolytic substrate specificity across protease classes (**Figure 4**). All MSP-MS cleavage sites were repressed in the presence of the serine protease inhibitor, suggesting that all P1↓P1’ pairs detected in the absence of inhibitors were related to serine protease activity. This also indicated that most exo-proteolytic activity in Lake Geneva is of serine protease origin. Cysteine and metalloprotease inhibitors partially removed cleavage specificity associated with arginine and lysine at positions P1 and P1’, revealing that these two amino acids are preferred for all three protease classes studied here. However, not all MSP-MS cleavage sites were repressed by these two inhibitors, suggesting that cysteine and metalloproteases are more restrictive in their substrate specificities. Notably, the P1↓P1’ cleavage frequency heatmaps in the presence of cysteine and metalloprotease inhibitors are highly similar. Whether the proteases of these two classes possess similar substrate specificities or whether the inhibition of one class affects the activity of proteases of the other remains to be investigated. Further work is additionally required to specifically identify the proteases present in lakewater.

The proteolytic fingerprints presented here offer a comprehensive overview of all microbial extracellular proteases present in Lake Geneva and reveal new insights into their functional capabilities. They add to the concept of enzymatic profiles for freshwater ecosystems, which have traditionally relied on exo-enzymatic activity measurements using fluorogenic probes such as LLßN^2,10,12,13^. These substrates are primarily cleaved by amino(exo)peptidases, mainly those with broad substrate specificities or those targeting the amino acid leucine^34^. Such activity measurements can represent 9.7% to 96% of total extracellular proteolytic activity levels in freshwater sources and may therefore greatly underestimate the contribution of endopeptidases^39^. Thus, the total extracellular proteolytic activities of the samples have not been fully reflected in most studies to date^39–42^. As the MSP-MS assay was designed to evaluate proteolytic cleavage by both exo- and endoproteases, this method captures a broader spectrum of extracellular proteases. This difference may explain the observed lack of correlation between the exo-proteolytic activity levels and the number of detected MSP-MS cleavage sites (**Supplementary Figure 2**). However, the MSP-MS assay also presents certain limitations. Notably the short length of the library peptides excludes secondary and tertiary structures which can be important for substrate recognition by some proteases. Nonetheless, this method provides a more comprehensive understanding of extracellular proteolytic activity and function.

Uncovering the proteolytic substrate specificities of microbial extracellular proteases is a key step towards better understanding protein degradation in lake ecosystems. This study describes the global proteolytic fingerprint of Lake Geneva and of two other Swiss lakes and provides insight into the protease classes present therein. While the predominant proteolytic substrate specificities were conserved over time, depth, and across lakes, variability was observed among minor substrate specificities. These findings help advance our understanding of protein degradation and may be used to predict the stability of peptidic compounds and contaminants in lakewater.

## 5. Conclusion

- The global proteolytic fingerprint of the extracellular proteases present in Lake Geneva was revealed using the MSP-MS assay. A clear preference for positively charged and certain non-polar amino acids either side of the cleavage site was observed, while cleavage of negatively charged and polar amino acids was disfavored.
- These patterns were conserved across seasons and water depth as well as across lakes of different trophic state, suggesting high conservation of microbial extracellular proteases and/or their substrate specificities.
- Our results present a global view of the functional capability of the extracellular proteases present in three Swiss lakes and provide new insights into protein degradation in lake ecosystems. These results may additionally be used to predict the stability and fate of peptidic contaminants in the environment.

## Author Contributions

**Josephine Meibom**: Conceptualization, Methodology, Investigation, Data Curation, Writing – Original Draft. **Natalie Wichmann**: Methodology, Writing – Review & Editing. **Michael Zumstein**: Resources, Writing – Review & Editing, Funding acquisition, Supervision. **Tamar Kohn**: Conceptualization, Writing – Review & Editing, Resources, Project administration, Funding acquisition.

## Acknowledgements

This work was funded by the Swiss National Science Foundation (grant no. 310030 215226). NW was funded by the Swiss National Science Foundation through the Ambizione Fellowship PZ00P2_193130 to MZ. We thank Professor Anthony O’Donoghue and Diego Trujillo (O’Donoghue laboratory, University of California San Diego) for providing information on and the R scripts for the MSP-MS assay data analysis. We thank L. Daniela Morales and Aina Astorch-Cardona for proving critical feedback on the manuscript.

## Declaration of Interests

We declare no known conflicts of interest.

## Data Availability

All data is available on Zenodo under link https://doi.org/10.5281/zenodo.15393523.

## Notes

https://doi.org/10.5281/zenodo.15393524

### Competing Interest Statement

The authors have declared no competing interest.

