## Supplemental figures and tables for "Proteolytic Activity and Substrate Specificity of Lake Geneva"

### Supporting Information

S1. Tables

| | Lake Geneva | Lake Neuchâtel | Lake Bret |
|---|---|---|---|
| **Surface Area** | 580.1 $km^2$ | 242 $km^2$ | 0.5 $km^2$ |
| **Volume** | 89 $km^3$ | 13.9 $km^3$ | 4.3 $km^3$ |
| **Maximum Depth** | 310 m | 153 m | 20 m |
| **Altitude** | 372 m | 429 m | 673 m |
| **Trophic State** | Mesotrophic | Oligotrophic | Eutrophic |
| **Catchment Size** | 7419 $km^2$ | 2462 $km^2$ | 23.3 $km^2$ |
| **Main Land Use in the Catchment Area** | Forests (44%) & Natural surfaces without vegetation (33%) | Agriculture (46%) & Forests (36%) | Agriculture (63%) & Forests (20%) |

**Supplementary Table 1**. Characteristics of Lakes Geneva, Neuchâtel, and Bret[1–3].

| | April '24 | July '24 | Sept. '24 | October '24 | February '25 |
|---|---|---|---|---|---|
| **April '24** | NA | 0.654 (0.002) | 0.558 (0.130) | 0.498 (0.130) | 0.427 (0.130) |
| **July '24** | 0.654 (0.002) | NA | 0.820 (3.67E-05) | 0.854 (3.67E-05) | 0.782 (3.67E-05) |
| **Sept. '24** | 0.558 (0.130) | 0.820 (3.67E-05) | NA | 0.838 (3.67E-05) | 0.850 (3.67E-05) |
| **October '24** | 0.498 (0.130) | 0.854 (3.67E-05) | 0.838 (3.67E-05) | NA | 0.867 (3.67E-05) |
| **February '25** | 0.427 (0.130) | 0.782 (3.67E-05) | 0.850 (3.67E-05) | 0.867 (3.67E-05) | NA |

**Supplementary Table 2**. Pearson Correlation Coefficients of amino acid frequencies surrounding the cleavage sites (P4-P4') in the detected MSP-MS cleavage products at t = 6h. Corrected p-values using the Benjamini-Hochberg method are indicated in parentheses.

S2. Figures

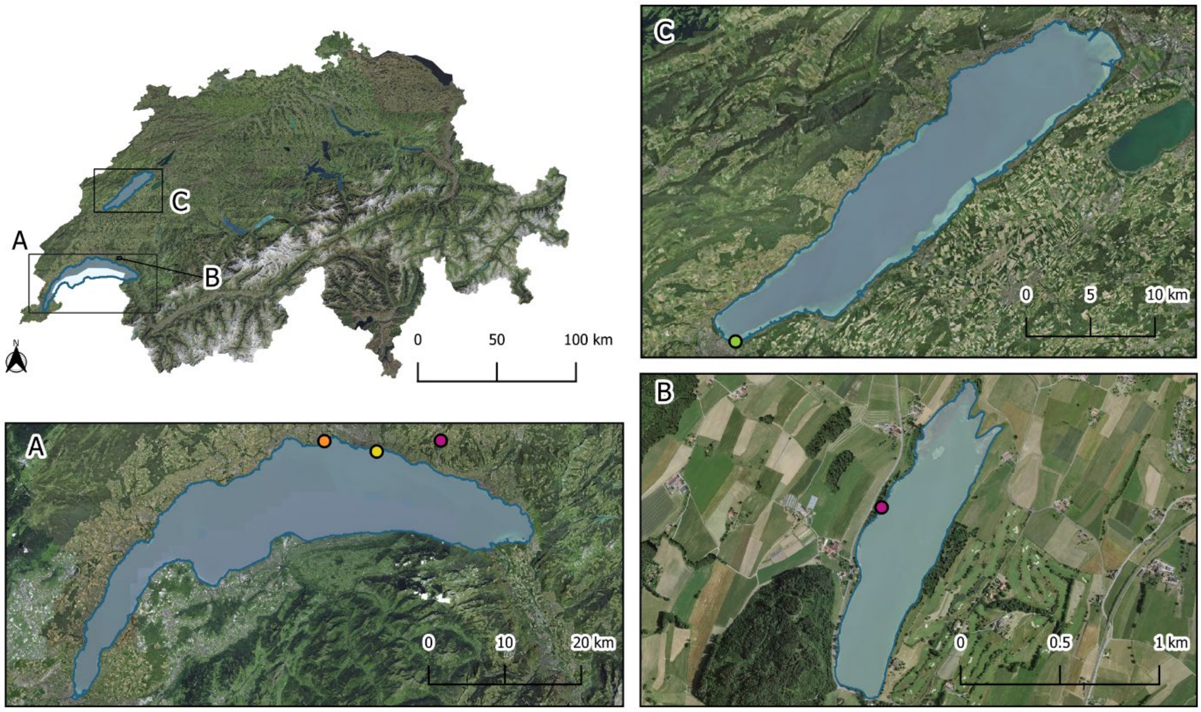


**Supplementary Figure 1**. Map locating Lake Geneva (**box A**), Lake Bret (**box B**), and Lake Neuchâtel (**box C**) in Switzerland. The sampling sites are indicated by colored dots: Saint-Sulpice (Lake Geneva, orange), LéXPLORE platform (Lake Geneva, yellow), shore of Lake Bret (pink), Yverdon-les-Bains (Lake Neuchâtel, green). The map was made using SWISSIMAGE 10 cm images provided by the Federal Office of Topography swisstopo with the QGIS software.

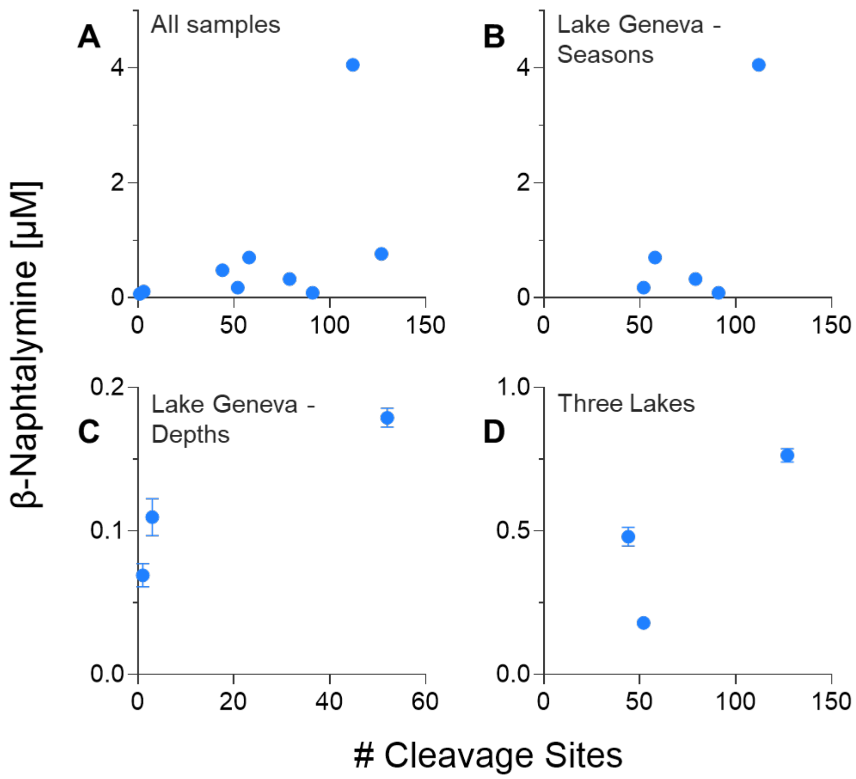


**Supplementary Figure 2**. Exo-proteolytic activity as measured by the L-leucine-ß-naphthylamide (LLßN) assay as a function of the number of the number of detected cleavage sites during incubation with the peptide library (t = 6h for all samples). **(A)** all samples. **(B)** Lake Geneva samples collected from April 2024 to February 2025, **(C)** Lake Geneva samples collected at different depths in October 2024, **(D)** samples collected from Lakes Geneva, Neuchâtel, and Bret in October 2024. Individual data points of triplicate measurements are presented with the geometric mean and standard deviation. Note the different scales.

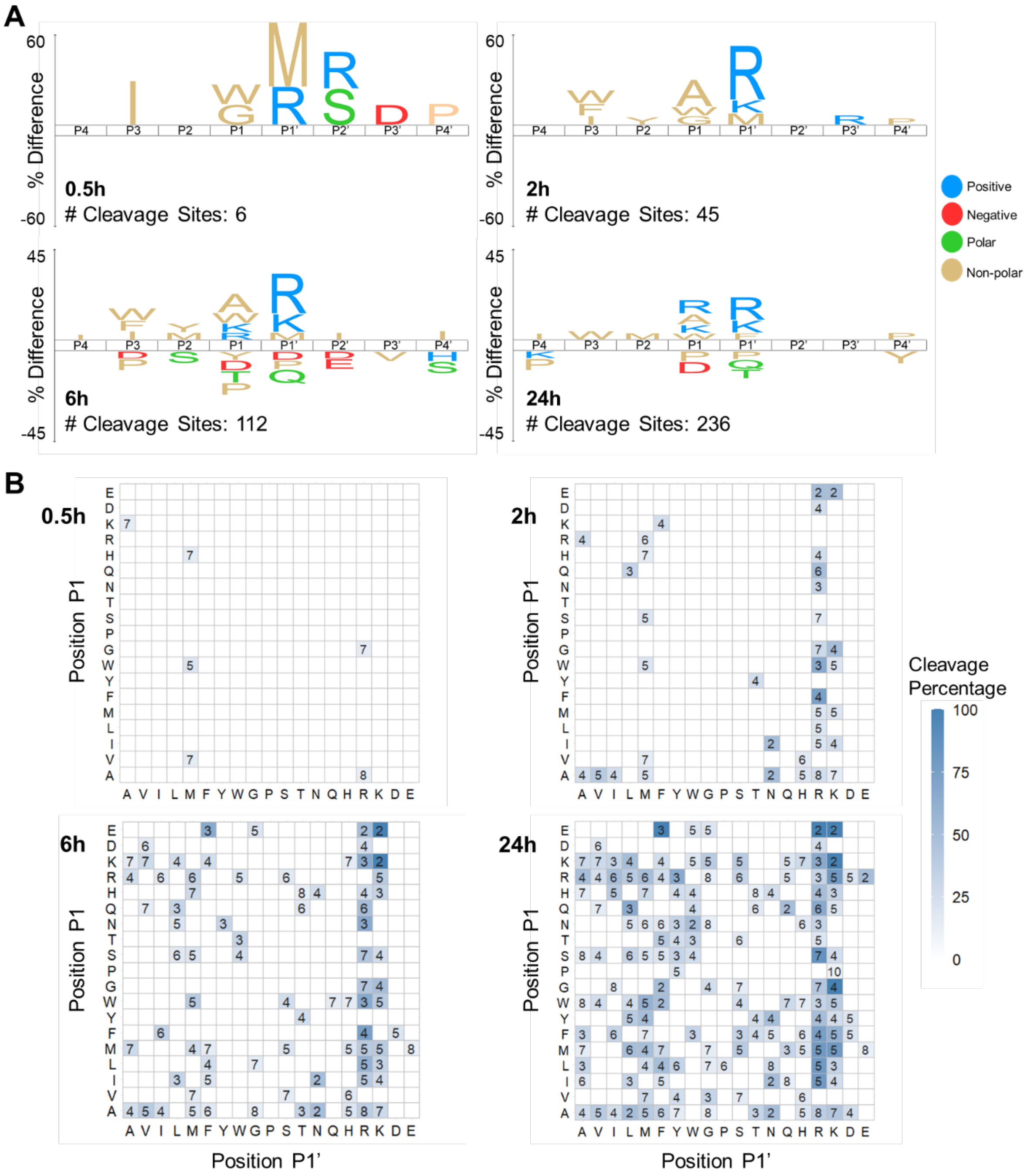


**Supplementary Figure 3**. Time course of the proteolytic fingerprint of Lake Geneva in April 2024. **(A)** iceLogo plots at t = 0.5h, 2h, 6h, and 24h. The number of detected cleavage sites is indicated for each timepoint. One-letter amino acid codes are used with M representing norleucine. Note the change in scale between timepoints. **(B)** P1↓P1' cleavage frequency heatmaps at t = 0.5h, 2h, 6h, and 24h. The number in each cell represents the number of occurrences of the corresponding P1↓P1' pair in the MSP-MS library. The color scale indicates the percentage of these that are cleaved during incubation in the sample.

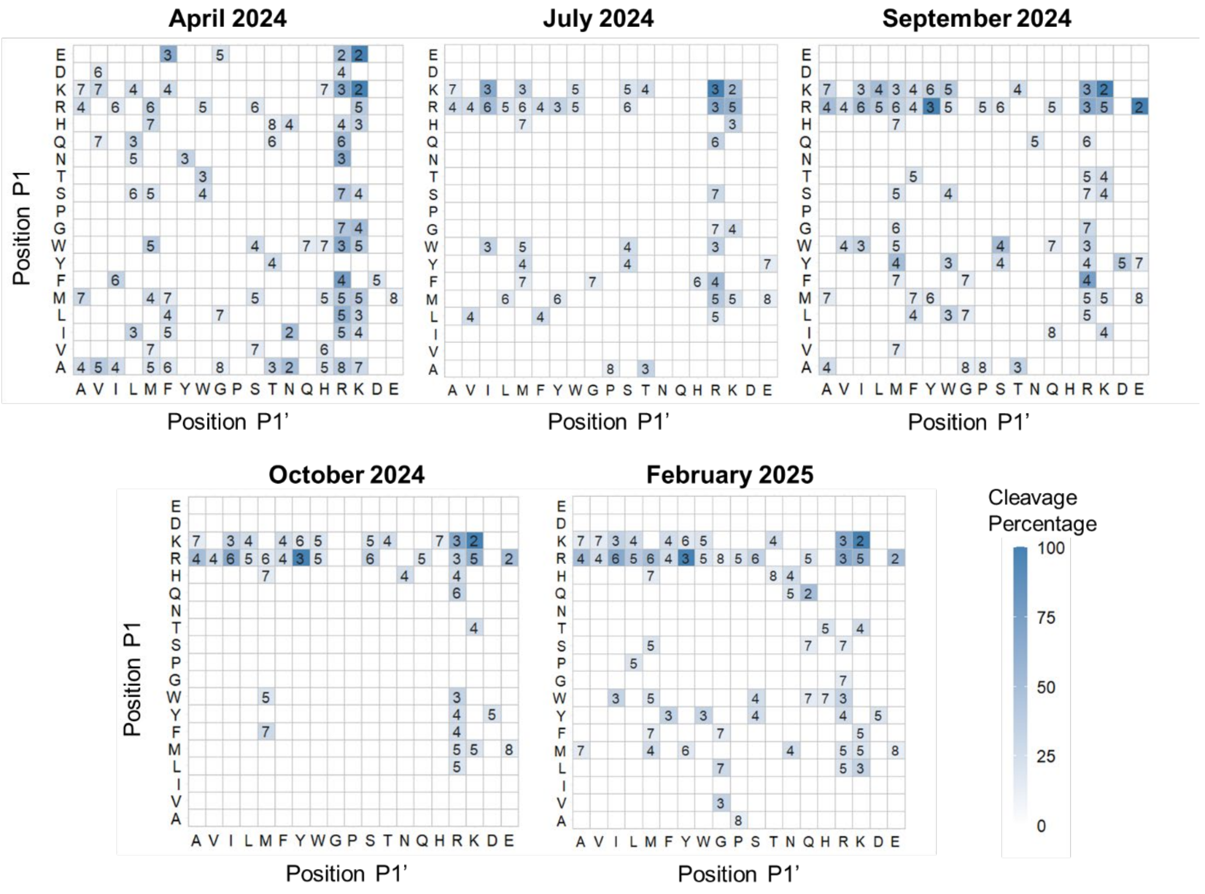


**Supplementary Figure 4.** P1↓P1’ cleavage frequency heatmaps of Lake Geneva at t = 6h for all seasons (April 2024, July 2024, September 2024, October 2024, and February 2025). The number in each cell represents the number of occurrences of the corresponding P1↓P1’ pair in the MSP-MS library. The color scale indicates the percentage of these that are cleaved during incubation in the sample.

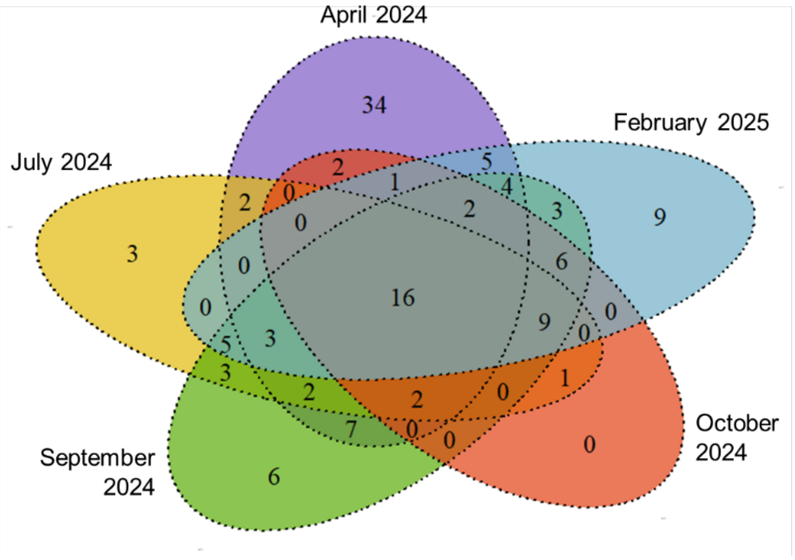


**Supplementary Figure 5**. Venn diagram representing the different P1↓P1' pairs cleaved in each lake (t = 6h). Each cleaved P1↓P1' pair is counted once, irrespective of its cleavage frequency in the sample.

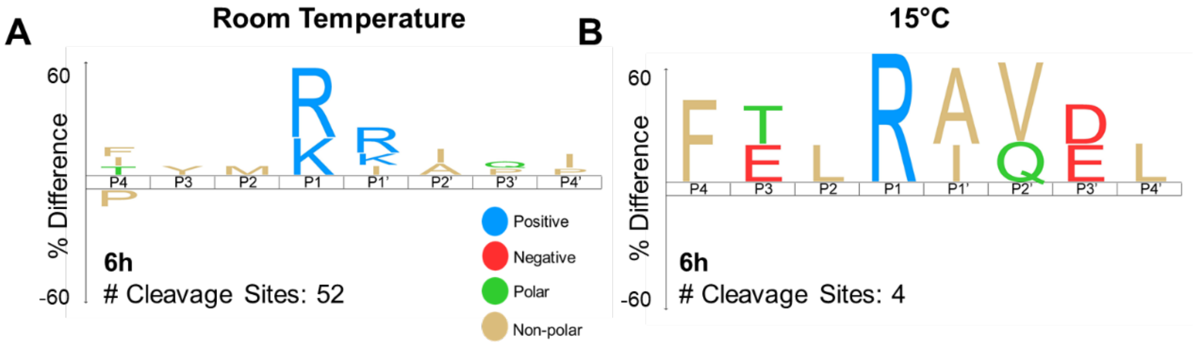


**Supplementary Figure 6**. Proteolytic fingerprint of Lake Geneva in October 2024 at t = 6h. The reactions were maintained at room temperature **(A)** or at 15°C **(B)**.

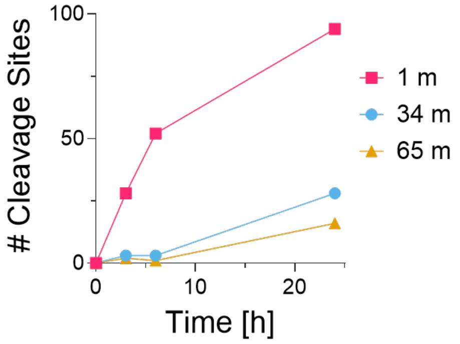


**Supplementary Figure 7**. The number of detected cleavage sites during incubation of the MSP-MS peptide library in Lake Geneva water collected in October 2024 at different depths (1 m, 34 m, and 65 m).

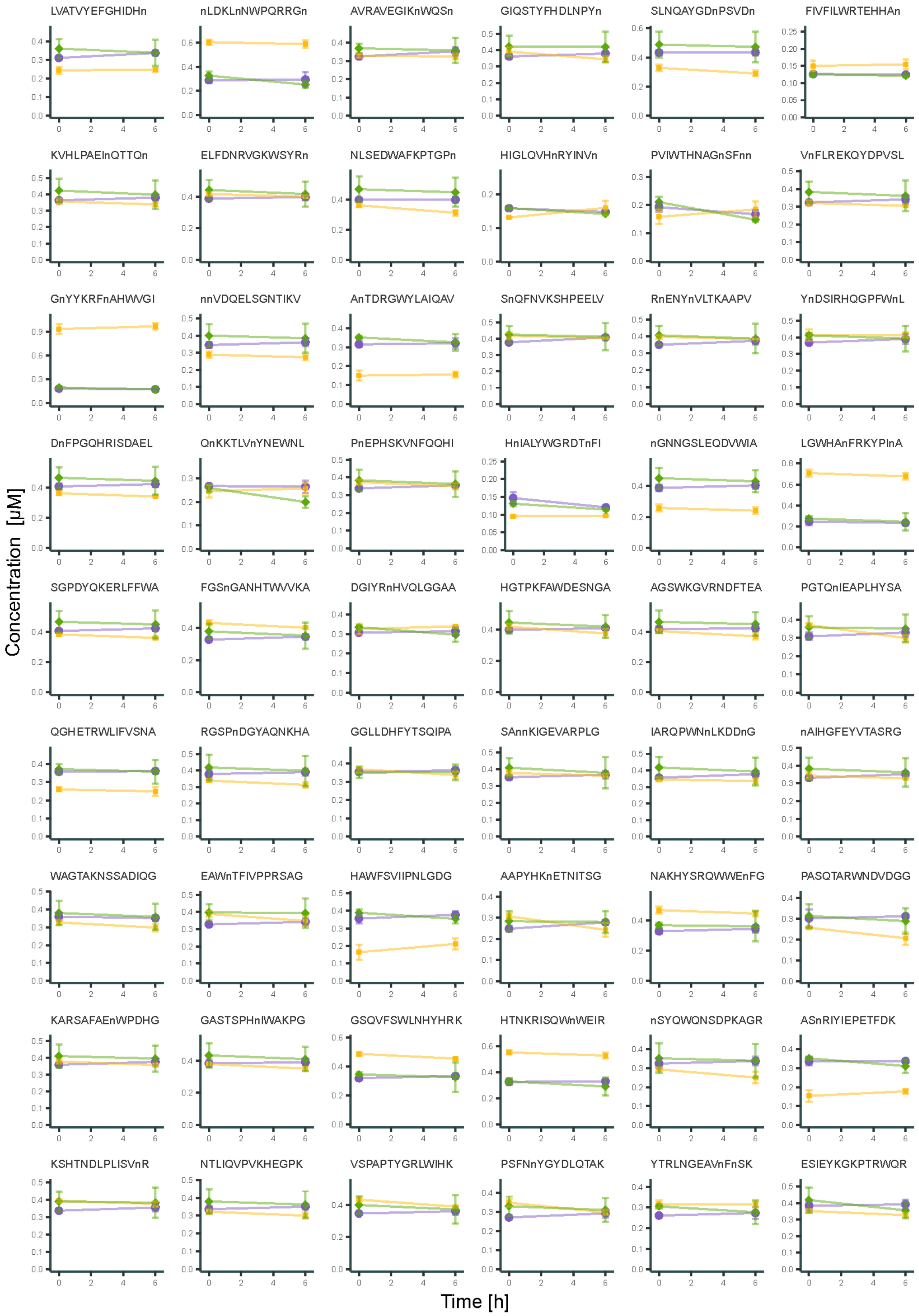

LVATVYEFGHIDHn
nLDKLnNWPQRRGn
AVRAVEGIKnWQSn
GIQSTYFHDLNPYn
SLNQAYGDnPSVDn
FIVFILWRTEHHAn
KVHLPAEInQTTQn
ELFDNRVGKWSYRn
NLSEDWAFKPTGPn
HIGLQVHnRYINVn
PVIWTHNAGnSFnn
VnFLREKQYDPVSL
GnYYKRFnAHWVGI
nnVDQELSGNTIKV
AnTDRGWYLAIQAV
SnQFNVKSHPEELV
RnENYnVLTKAAPV
YnDSIRHQGPFWnL
DnFPGQHRISDAEL
QnKKTLVnYNEWNL
PnEPHSKVNFQQHI
HnIALYWGRDTnFI
nGNNGSLEQDVWIA
LGWHAnFRKYPInA
SGPDYQKERLFFWA
FGSnGANHTWVVKA
DGIYRnHVQLGGAA
HGTPKFAWDESNGA
AGSWKGVRNDFTEA
PGTQnIEAPLHYSA
QGHETRWLIFVSNA
RGSPnDGYAQNKHA
GGLLDHFYTSQIPA
SAnnKIGEVARPLG
IARQPWNnLKDDnG
nAIHGFEYVTASRG
WAGTAKNSSADIQG
EAWnTFIVPPRSAG
HAWFSVIIPNLGDG
AAPYHKnETNITSG
NAKHYSRQWWEnFG
PASQTARWNDVDGG
KARSAFAEnWPDHG
GASTSPHnIWAKPG
GSQVFSWLNHYHRK
HTNKRISQWnWEIR
nSYQWQNSDPKAGR
ASnRIYIEPETFDK
KSHTNDLPLISVnR
NTLIQVPVKHEGPK
VSPAPTYGRLWIHK
PSFNnYGYDLQTAK
YTRLNGEAVnFnSK
ESIEYKGKPTRWQR
Concentration [µM]
Time [h]

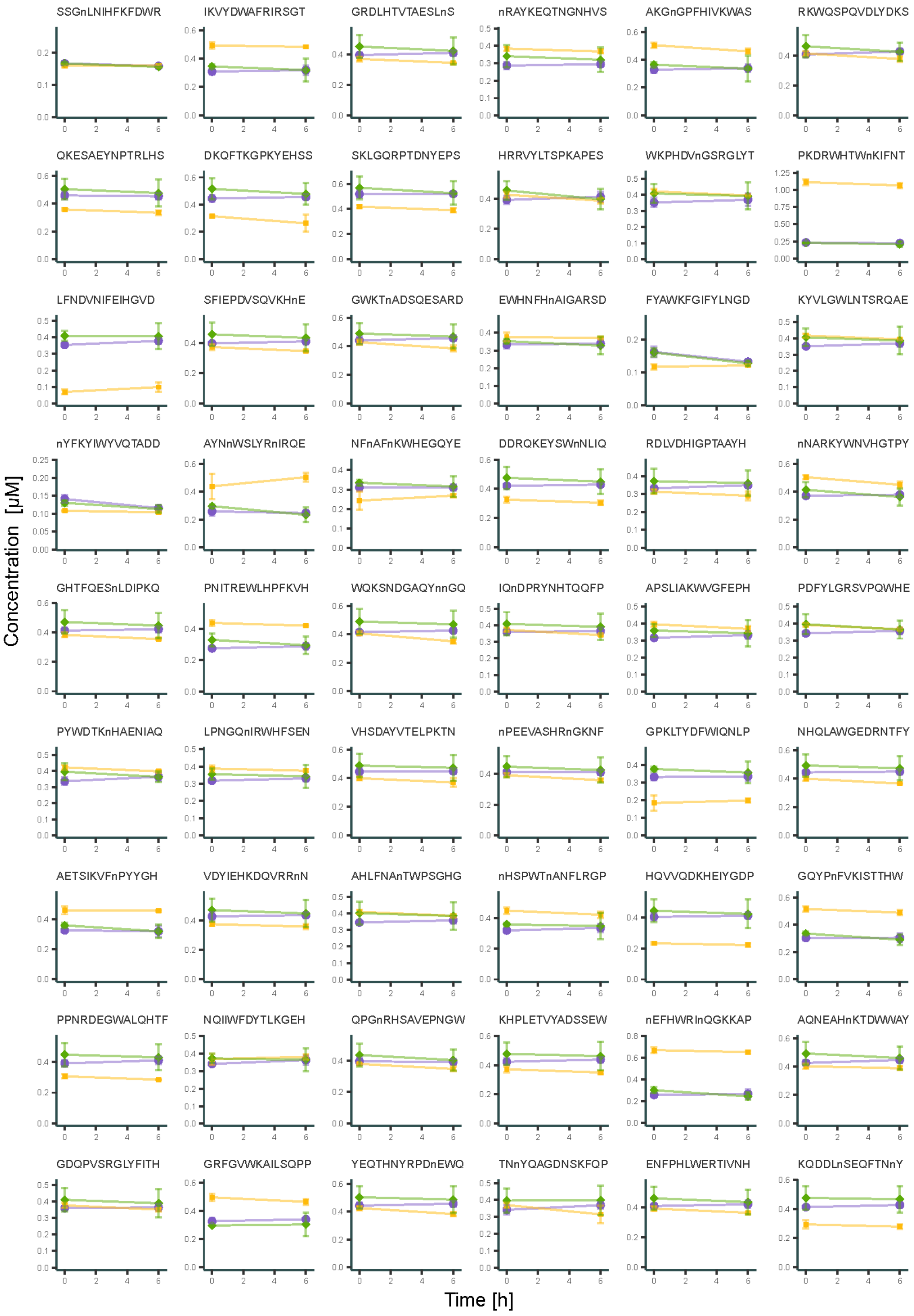
SSGnLNIHFKFDWR
IKVYDWAFRIRSGT
GRDLHTVTAESLnS
nRAYKEQTNGNHVS
AKGnGPFHIVKWAS
RKWQSPQVDLYDKS
QKESAEYNPTRLHS
DKQFTKGPKYEHSS
SKLGQRPTDNYEPS
HRRVYLTSPKAPES
WKPHDVnGSRGLYT
PKDRWHTWnKIFNT
LFNDVNIFEIHGVD
SFIEPDVSQVKHnE
GWKTnADSQESARD
EWHNFHnAIGARSD
FYAWKFGIFYLNGD
KYVLGWLNTSRQAE
nYFKYIWYVQTADD
AYNnWSLYRnIRQE
NFnAFnKWHEGQYE
DDRQKEYSWnNLIQ
RDLVDHIGPTAAYH
nNARKYWNVHGTPY
GHTFQESnLDIPKQ
PNITREWLHPFKVH
WQKSNDGAQYnnGQ
IQnDPRYNHTQQFP
APSLIAKWVGFEPH
PDFYLGRSVPQWHE
PYWDTKnHAENIAQ
LPNGQnIRWHFSEN
VHSDAYVTELPKTN
nPEEVASHRnGKNF
GPKLTYDFWIQNLP
NHQLAWGEDRNTFY
AETSIKVFnPYYGH
VDYIEHKDQVRRnN
AHLFNAnTWPSGHG
nHSPWTnANFLRGP
HQVVQDKHEIYGDP
GQYPnFVKISTTHW
PPNRDEGWALQHTF
NQIIWFDYTLKGEH
QPGnRHSAVEPNGW
KHPLETVYADSSEW
nEFHWRInQGKKAP
AQNEAHnKTDWWAY
GDQPVSRGLYFITH
GRFGVWKAILSQPP
YEQTHNYRPDnEWQ
TNnYQAGDNSKFQP
ENFPHLWERTIVNH
KQDDLnSEQFTNnY
Concentration [μM]
Time [h]

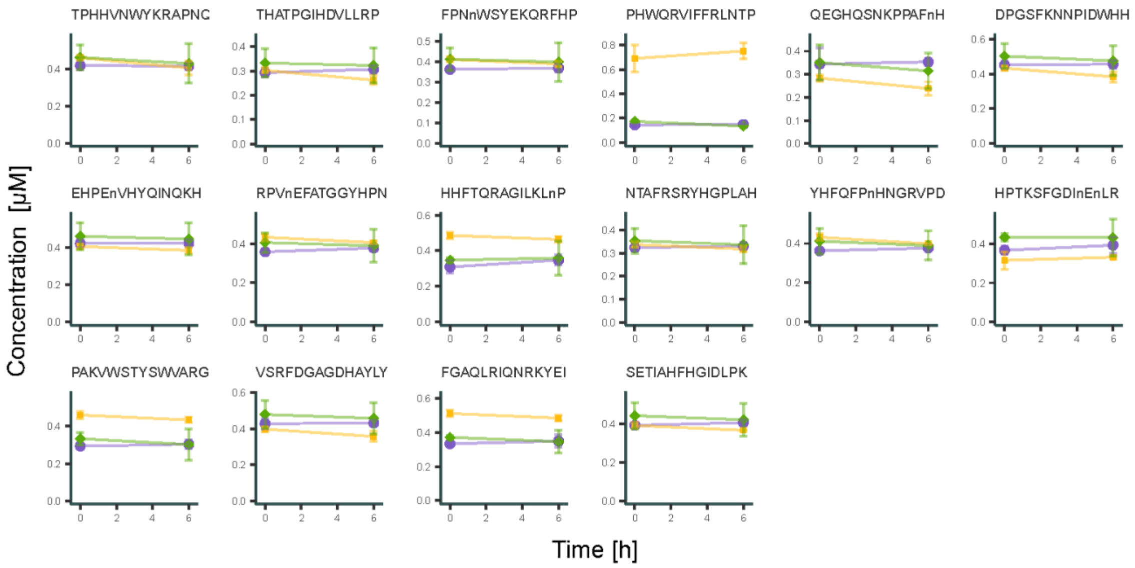


**Supplementary Figure 8**. Stability curves of the 124 MSP-MS library peptides in sterile lakewater (July 2024) in the absence and presence of class-specific protease inhibitors. No inhibitor: purple dots. Serine protease inhibitor: yellow squares. Metalloprotease inhibitor: green diamonds. The mean and standard deviation of triplicate values are presented. Phenylmethanesulfonyl fluoride (PMSF) was used as the serine protease inhibitor and GM6001 was used as the metalloprotease inhibitor.

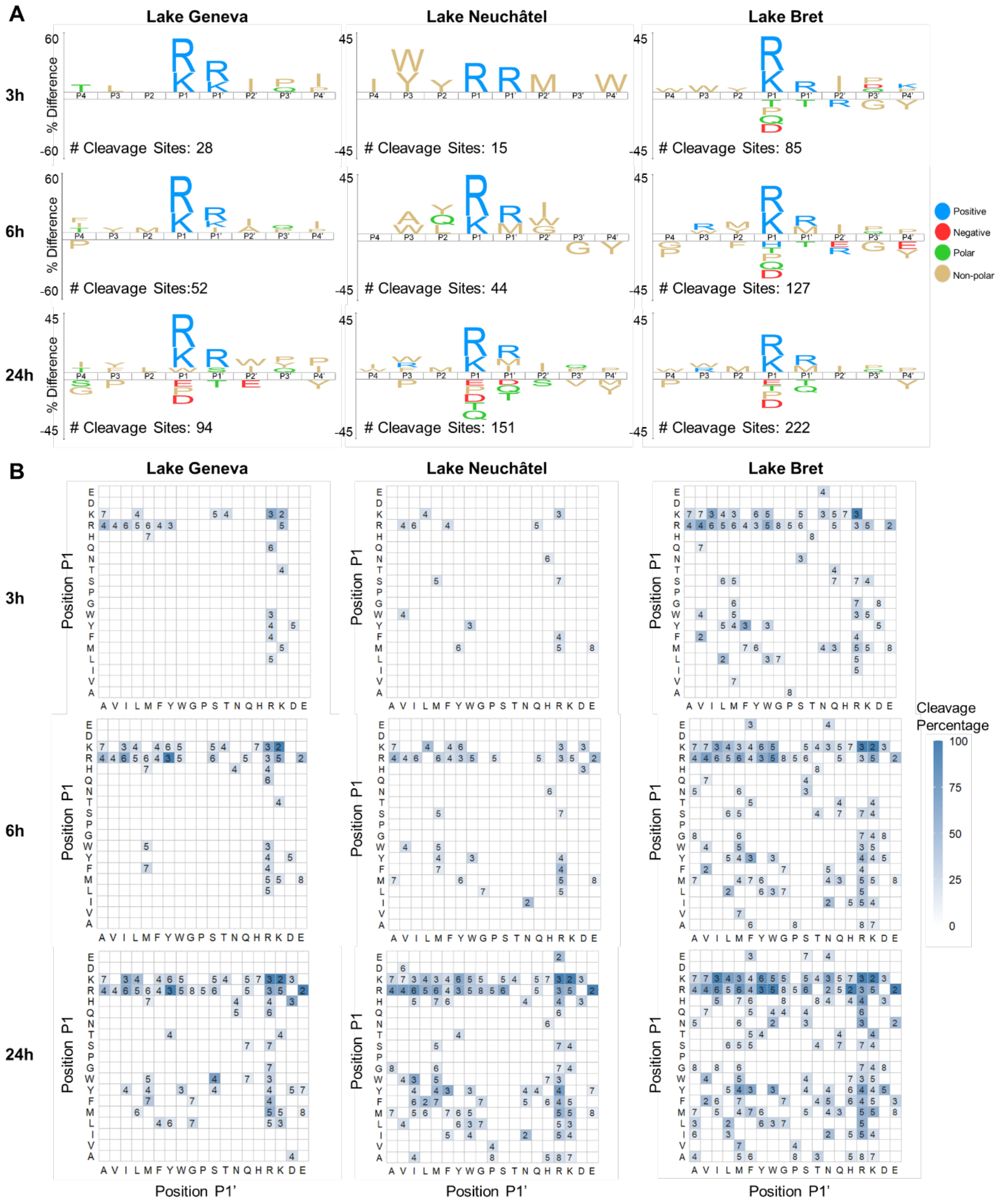


**Supplementary Figure 9**. Time course of the proteolytic fingerprint of Lakes Geneva, Neuchâtel, and Bret in October 2024. **(A)** iceLogo plots at t = 3h, 6h, and 24h. The number of detected cleavage sites is indicated for each timepoint. One-letter amino acid codes are used with M representing norleucine. Note the change in scale between timepoints. **(B)** P1↓P1' cleavage frequency heatmaps at t = 3h, 6h, and 24h. The number in each cell represents the number of occurrences of the corresponding P1↓P1' pair in the MSP-MS library. The color scale indicates the percentage of these that are cleaved during incubation in the sample.
